# A Cost-effective, High-throughput, Highly Accurate Genotyping Method for Outbred Populations

**DOI:** 10.1101/2024.07.17.603984

**Authors:** Denghui Chen, Apurva S. Chitre, Khai-Minh H. Nguyen, Katarina Cohen, Beverly Peng, Kendra S. Ziegler, Faith Okamoto, Bonnie Lin, Benjamin B. Johnson, Thiago M. Sanches, Riyan Cheng, Oksana Polesskaya, Abraham A. Palmer

## Abstract

Affordable sequencing and genotyping methods are essential for large scale genome-wide association studies. While genotyping microarrays and reference panels for imputation are available for human subjects, non-human model systems often lack such options. Our lab previously demonstrated an efficient and cost-effective method to genotype heterogeneous stock rats using double-digest genotyping-by-sequencing. However, low-coverage whole-genome sequencing offers an alternative method that has several advantages. Here, we describe a cost-effective, high-throughput, high-accuracy genotyping method for N/NIH heterogeneous stock rats that can use a combination of sequencing data previously generated by double-digest genotyping-by-sequencing and more recently generated by low-coverage whole-genome-sequencing data. Using double-digest genotyping-by-sequencing data from 5,745 heterogeneous stock rats (mean 0.21x coverage) and low-coverage whole-genome-sequencing data from 8,760 heterogeneous stock rats (mean 0.27x coverage), we can impute 7.32 million bi-allelic single-nucleotide polymorphisms with a concordance rate >99.76% compared to high-coverage (mean 33.26x coverage) whole-genome sequencing data for a subset of the same individuals. Our results demonstrate the feasibility of using sequencing data from double-digest genotyping-by-sequencing or low-coverage whole-genome-sequencing for accurate genotyping, and demonstrate techniques that may also be useful for other genetic studies in non-human subjects.

**Article Summary:** Heterogeneous stock rats were derived by interbreeding eight inbred founders in 1984, and have been maintained as an outbred population for more than 100 generations. Heterogeneous stock rats offer a high degree of genetic and phenotypic diversity, and have been extensively used for genetic studies. Here, we describe a cost-effective and high-throughput genotyping method for heterogeneous stock rats. We applied the method to 15,552 heterogeneous stock rats. The resulting genotypes were highly accurate. These techniques may be useful for genetic studies in other non-human subjects.

## Introduction

In both humans and model organisms, genome-wide association studies (**GWAS**) are valuable for identifying genetic variants associated with diseases and other complex traits. GWAS results facilitate the discovery of novel biological pathways and potential therapeutic targets (Palmer et al. 2021; Uffelmann et al. 2021; Alliance of Genome Resources Consortium 2022; Abdellaoui et al. 2023). The success of large scale population and quantitative genetics studies depends on the availability of dense and high quality genotype data (Welter et al. 2014). Single-nucleotide polymorphism (**SNP**) arrays, paired with reference panels (e.g. HapMap or the 1000 Genomes Project), are commonly used to infer genotypes and perform genetic studies in humans (Frazer et al. 2007; Marchini and Howie 2010; McVean et al. 2012; Uffelmann et al. 2021; Aganezov et al. 2022). However, SNP arrays often perform poorly when applied to populations other than the one used for array design, leading to a need for costly development of population-specific SNP arrays (Didion et al. 2012). This issue is even more critical in model organisms, where population structure is often very pronounced (Gileta et al. 2022). An alternative to genotyping microarrays is to use next-generation sequencing. Because sequencing at sufficient depth to make calls directly remains expensive, low coverage sequencing paired with imputation from reference panels provides a more economical solution (Davies et al. 2016; Petter et al. 2020; Li et al. 2021; Wasik et al. 2021; Li et al. 2024).

Our lab has performed GWAS using various mouse and rat populations (Chitre et al. 2020; Zhou et al. 2020; Gileta et al. 2022; Gunturkun et al. 2022; Parker et al. 2022; Chitre et al. 2023; Fowler et al. 2023). In particular, we have now phenotyped and genotyped almost 20,000 N/NIH heterogeneous stock (**HS**) rats. HS rats were created in 1984 by intercrossing eight inbred rat strains (ACI/N, BN/SsN, BUF/N, F344/N, M520/N, MR/N, WKY/N, and WN/N). To genotype outbred mice and rats we have used genotyping-by-sequencing (**GBS**) (Elshire et al. 2011; Parker et al. 2016; Gonzales et al. 2018) and subsequently double-digest genotyping-by-sequencing (**ddGBS**) protocols, followed by imputation (Gileta et al. 2020). More recently, we have reported on our use of commercial whole-genome-sequencing (**WGS**) library preparation kits to generate low-coverage WGS (**lcWGS**) data, followed by imputation using outbred mice (Davies et al. 2016; Nicod et al. 2016; Zou et al. 2022). However, we have not previously reported on our methods for genotyping rats using lcWGS followed by imputation, nor have we reported a method for jointly calling genotypes using a combination of ddGBS and lcWGS data.

In this paper, we present a cost-effective, high-throughput, and highly accurate genotyping method for HS rats that utilizes both previously-generated ddGBS data and more recently generated lcWGS data. This method allowed us to impute 7.32 million bi-allelic SNPs with a concordance rate of >99.76% compared to genotypes obtained from 33.26x coverage WGS without imputation for a subset of the same individuals.

## Materials and Methods

### Animals

As reviewed elsewhere, the N/NIH HS rat population was created by interbreeding eight inbred rat strains (ACI/N, BN/SsN, BUF/N, F344/N, M520/N, MR/N, WKY/N, and WN/N) in the mid 1980s (Solberg Woods and Palmer 2019). Since then, HS rats have been maintained as an outbred population for more than 100 generations. Because they have been maintained as an outbred population for such a long time, HS rats possess short haplotypes that are derived from the eight inbred founders, making them ideal for high-resolution genetic mapping (Johannesson et al. 2009; Baud et al. 2013; Woods and Mott 2017; Solberg Woods and Palmer 2019). In this study, we used sequence data from a total of 15,552 HS rats (7,797 males, 7,755 females) from generation 81 to 97 that were bred at the Medical College of Wisconsin (RRID: RGD_2314009), Wake Forest University (RRID: RGD_13673907), the University of California San Diego (RRID: RGD_155269102), the University of Tennessee Health Sciences Center, or Oregon Health and Sciences University. All procedures that occurred prior to tissue collection were approved by the relevant Institutional Animal Care and Use Committees. As described in the following sections, of the 15,552 HS rats, 477 were sequenced with both ddGBS and lcWGS. 88 of those 477 were also whole genome sequenced at an average depth of 33.26x; we refer to those 88 rats as the ‘truth set’.

### ddGBS Sequencing

Of the 15,552 HS rats used in this study, 6,379 individuals (3,219 males; 3,160 females) were sequenced using a ddGBS library preparation protocol described by (Gileta et al. 2020). Briefly, DNA was extracted from spleen tissues using Agencourt DNAdvance Kit (Beckman Coulter Life Sciences, Indianapolis, IN) and digested using the restriction enzymes Pstl and NlaIII. After adapter ligation, DNA purification and library pooling, sample DNA was sequenced as 48 samples per library on Illumina HiSeq 4000 with 100 bp single-end reads at the University of California San Diego Institute for Genomic Medicine Genomics Center (**UCSD IGM**).

### lcWGS Sequencing

In addition, 9,173 (4,578 males; 4,595 females) of 15,552 HS rats underwent lcWGS sequencing. DNA was extracted from spleen tissues using the Agencourt DNAdvance Kit, and the Twist 96-Plex Library Prep Kit (Twist Bioscience, South San Francisco, CA) was used for library preparation following the manufacturer’s protocol. The samples’ DNA were sequenced as 96 samples per library on Illumina NovaSeq 4000 or 6000 with 150 bp paired-end reads at UCSD IGM. DNA extraction, normalization, randomization and library preparation were all performed on the EPmotion 5075 (Eppendorf, Hamburg, Germany) liquid handling robot. Detailed lcWGS protocols for many of these steps can be found in the Center for GWAS in Outbred Rats Database protocol repository on protocols.io (https://www.protocols.io/workspaces/cgord, spleen cutting: http://dx.doi.org/10.17504/protocols.io.36wgq7nryvk5/v1, DNA extraction: http://dx.doi.org/10.17504/protocols.io.8epv59reng1b/v1, normalization and randomization: http://dx.doi.org/10.17504/protocols.io.261genw5dg47/v1, library preparation: http://dx.doi.org/10.17504/protocols.io.j8nlkkm85l5r/v1, pooling and sequencing: http://dx.doi.org/10.17504/protocols.io.yxmvmnw29g3p/v1).

### Reference Panel Preparation

To obtain the best possible imputation reference panel for outbred HS rats, we used consensus bi-allelic homozygous SNP calls from three different inbred HS rats founder datasets. The first dataset was produced from publicly available 30.34x coverage WGS sequences (NCBI SRA: PRJNA487943) using the Genome Analysis Toolkit (**GATK**) joint calling workflow (Method S2 and Figure S1 in Supplementary Material) (Ramdas et al. 2019; Van der Auwera and O’Connor 2020). In that dataset, BN/SsN and MR/N are female, and other rats are male. The second dataset was produced using the same GATK joint calling workflow using an independent dataset with an average of 41.81x coverage WGS sequences (NCBI SRA: PRJNA1048943) generated with high coverage WGS sequencing procedures (Method S1, Method S2 and Figure S1 in Supplementary Material). Details of this dataset have not been previously published. In this dataset, all eight HS founders were male. The third dataset was produced using the same 41.81x coverage WGS sequences, but using the DeepVariant multi-sample calling workflow (Method S3 and Figure S2 in Supplementary Material). For autosomal chromosomes, chromosome X and mitochondria, 7,406,667, 184,934 and 117 SNPs respectively that had consensus homozygous genotypes across all three callsets were retained, however, because BN/SsN and MR/N in the first dataset are female, we dropped them from the consensus check process for chromosome Y, resulting in 5,220 consensus homozygous SNPs for chromosome Y. In total, 7,596,938 SNPs were retained for the reference panel.

### Bi-allelic SNP Positions Preparation

We employed STITCH for the imputation process. STITCH was designed for imputing bi-allelic SNPs in lcWGS reads by constructing haplotypes (Davies et al. 2016). STITCH accepts a position file for the bi-allelic SNPs to be imputed. In order to capture the common variants derived from the HS founders, as well as new SNPs observed in recent generations of the outbred HS population, we compiled the SNP position file using bi-allelic SNPs discovered in the founder datasets mentioned above and in 88 HS rats (44 males; 44 females). Variants in the subset of 88 HS rats were called on 33.26x coverage WGS sequences (NCBI SRA: PRJNA1076141) using the GATK joint calling workflow (Method S1, Method S2 and Figure S1 in Supplementary Material). The resulting SNPs position file contained 10,684,883 SNPs with 10,227,209 on autosomal chromosomes, 331,389 on chromosome X, 126,141 on chromosome Y and 144 on mitochondria.

### Truth Set Preparation

To assess the quality of imputed genotypes, we sequenced the aforementioned 88 HS outbred rats using three methods: ddGBS, lcWGS and high coverage WGS (33.26x). The bi-allelic SNPs imputed from the ddGBS and lcWGS genotyping pipeline were compared with the variants discovered on high coverage WGS GATK joint calling pipeline (Method S1, Method S2 and Figure S1 in Supplementary Material). We treated the genotypes called by high coverage WGS as our truth set, and used them to check the concordance of the genotypes imputed with other two methods.

### Genotyping

Our full bioinformatic pipeline is outlined in Figure 1. The pipeline inputs each sample’s raw ddGBS or lcWGS sequences, maps them to Rattus norvegicus reference genome mRatBN7.2 (NCBI Genome Assembly Accession: GCF_015227675.2) in parallel, and then jointly imputes bi-allelic SNPs. The complete source code for the pipeline can be found in the Palmer Lab GitHub repository (https://github.com/Palmer-Lab-UCSD/HS-Rats-Genotyping-Pipeline, DOI:https://doi.org/10.5281/zenodo.10002191).

**Figure 1.**
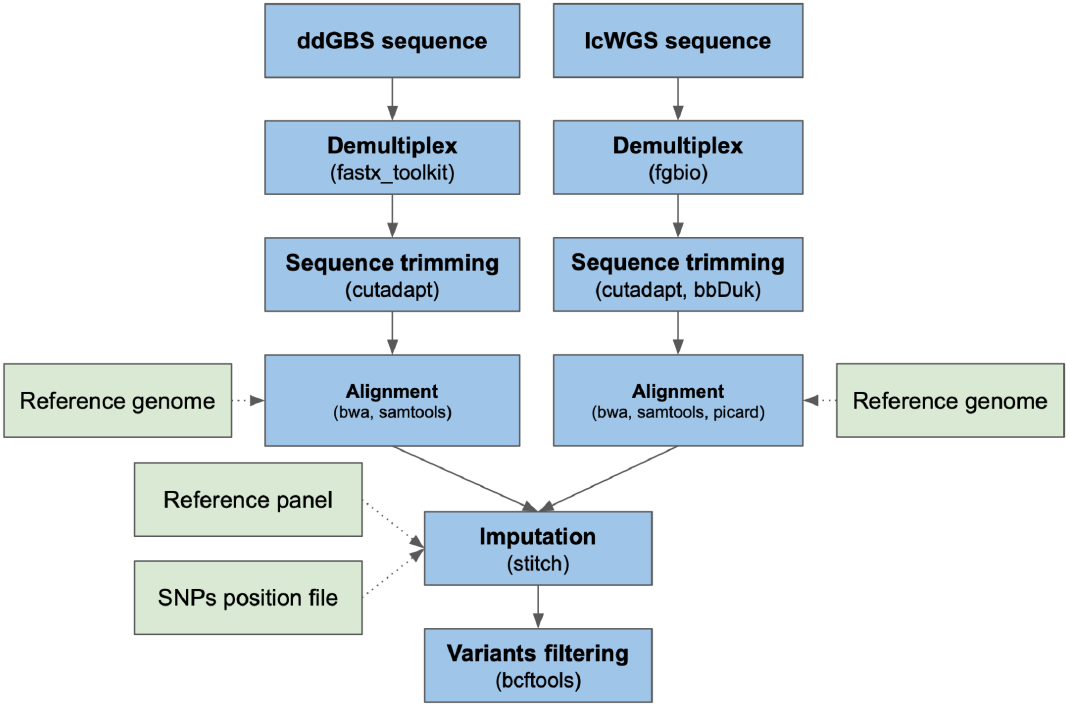
Genotyping pipeline flow chart.

ddGBS sequences were demultiplexed using fastx_toolkit v0.0.14 (Hannon Lab 2010). Barcode, adapter and quality trimming were subsequently performed using Cutadapt v4.1 (Martin 2011) with 25 bp as the minimum length per read and 20 as the minimum base quality. BWA-mem v0.7.17 (Li 2013) was used to align ddGBS sequences with a constraint of an alignment score greater than 20, and the aligned BAM files were sorted and indexed by coordinates using SAMtools v1.14 (Danecek et al. 2021) for fast random access.

lcWGS sequences were demultiplexed using fgbio v1.3.0 (Tim and Nils 2023). BBDuk v38.94 (Bushnell) (ktrim=r, k=23, mink=11, hdist=1, trimpolyg=50, tpe, tbo) was used to trim adapters, and Cutadapt v4.1 (Martin 2011) was used to trim sequences with Phred base quality less than 5 and length shorter than 70 bp. Alignment of the lcWGS sequences was carried out using BWA-mem v0.7.17 (Li 2013). Duplicated reads were marked using Picard v2.25.7 (Broad Institute 2019) and indexed by coordinates using SAMtools v1.14 (Danecek et al. 2021) for fast random access.

Aligned sequences were used to jointly impute bi-allelic SNPs at given positions with STITCH v1.6.6 (Davies et al. 2016) (niterations=2, k=8, nGen=100). During the imputation step, a reference panel based on the genotypes of the 8 inbred founder strains and the SNPs position file mentioned above were provided to STITCH to construct haplotypes for imputation. To increase computational efficiency, imputation was performed parallelly on chromosome chunks with a one megabase buffer on each end. Each chunk had a length of at least seven megabases and contained at least one thousand SNPs. Then, we used BCFtools v1.14 (Danecek et al. 2021) to concatenate the chunks back to individual chromosomes.

### SNPs Quality Control

Following the imputation process, we implemented a quality control procedure to filter out SNPs with low genotype quality. A total of 10,684,883 bi-allelic SNPs were imputed using our genotyping pipeline. Among them, we removed 2,737,742 SNPs with an imputation info score less than 0.9 using BCFtools v1.14 (Danecek et al. 2021). Furthermore, we filtered out 623,881 SNPs that have low concordance with the ground truth dataset described above. As a result, we retained 7,323,260 SNPs. The genotypes after quality control can be found in UC San Diego Library Digital Collections https://doi.org/10.6075/J0445MPC.

### Sample Quality Control

A sample quality control step was also performed to ensure sample quality. In total, 15,552 samples, representing 14,629 unique outbred HS rats, were used in this study. We excluded 66 samples whose ratio of mapped reads on chromosome X and Y were incompatible with their reported sex (Figure S3 in Supplementary Material). We also excluded 153 samples with a genotype missing rate exceeding 0.1 or a genotype heterozygosity rate falling outside the range of ± 4 standard deviations (Figure S4 in Supplementary Material). Because of the differences between ddGBS and lcWGS data, we conducted these two sample quality control criteria for different sequencing methods separately. Additionally, in the cases where we had multiple sequencing runs for the same samples, we kept only the one with the highest number of sequence reads. This quality control process resulted in the retention of 14,505 distinct HS rats (7,283 males, 7,222 females) with 5,745 individuals from ddGBS (2,903 males; 2,842 females) and 8,760 individuals from lcWGS (4,380 males; 4,380 females).

## Results

### Sequence Statistics

Our genotyping pipeline was applied to 15,552 samples, representing 14,629 unique outbred HS rats. 14,505 distinct samples were retained after the quality control steps described in the methods section, 5,745 of which were sequenced using ddGBS and 8,760 using lcWGS.

After demultiplexing and aligning to reference genome mRatBN7.2 (NCBI Genome Assembly Accession: GCF_015227675.2), a mean of 8.44 million 100 bp reads per sample were mapped to the reference genome in the case of ddGBS (Figure 2A). Because of the double restriction enzyme digestion employed in ddGBS, only the chromosomal regions near the enzyme cut sites were sequenced. This led to ddGBS sequences covering 4.97% of the genome per sample, with a mean coverage of 4.22x at each captured site (Figure 2B, 2C). Consequently, this approach resulted in an average mapped coverage of 0.21x per sample across the entire genome although that coverage was highly non uniform, by design (Figure 2D).

**Figure 2.**
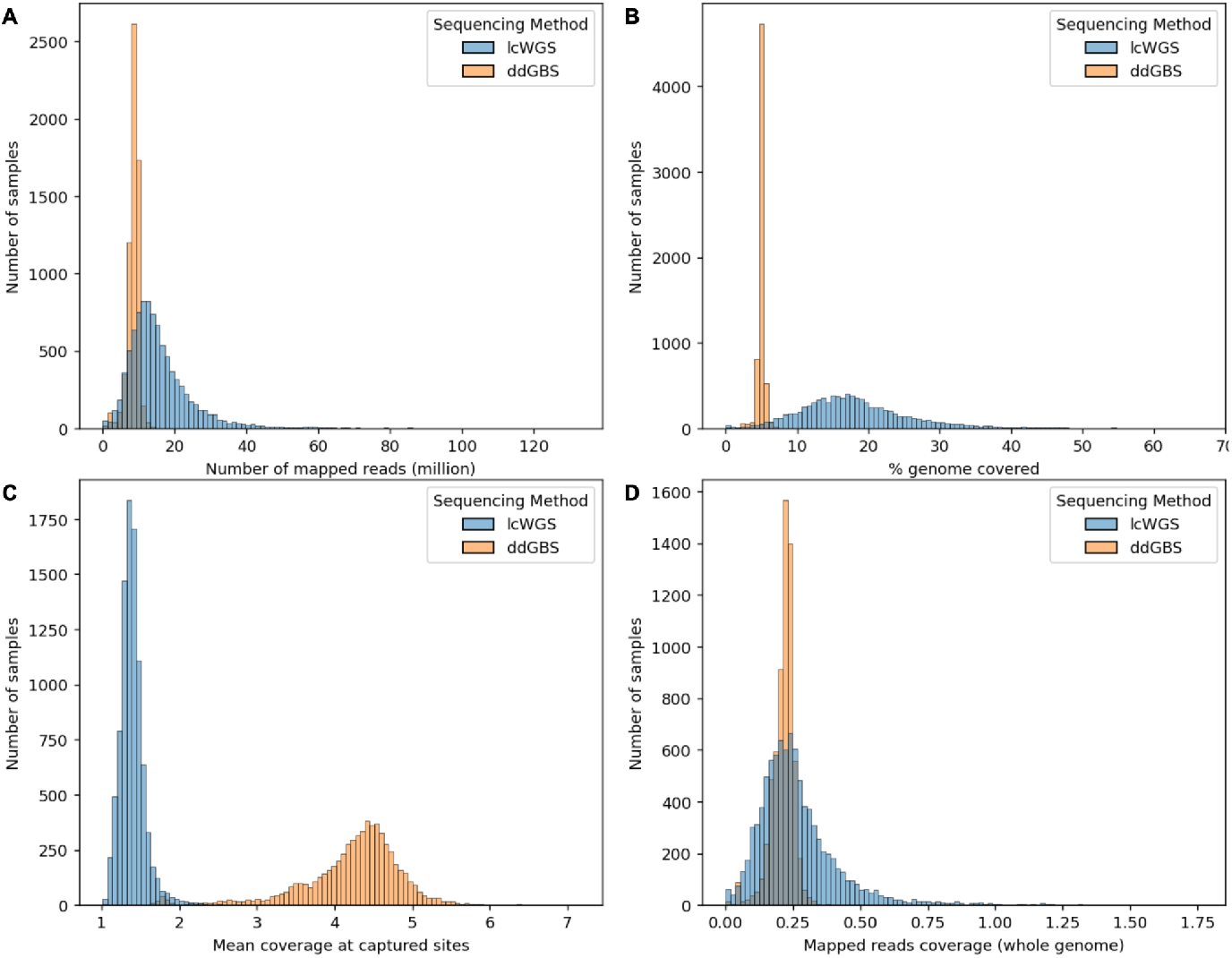
Aligned sequence statistics. **A**. Number of reads mapped to reference genome (million). ddGBS mean: 8.44, SD: 1.65; lcWGS mean: 16.03, SD: 10.32. **B**. Percentage of genome covered by mapped reads in width (%). ddGBS mean: 4.97, SD: 0.54; lcWGS mean: 18.28, SD: 8.31. **C**. Mean coverage at captured sites. ddGBS mean: 4.22x, SD: 0.67x; lcWGS mean: 1.39x, SD: 0.16x. **D**. Mapped reads coverage genome-wide. ddGBS mean: 0.21x, SD: 0.04x; lcWGS mean: 0.27x, SD: 0.16x.

For lcWGS, a mean of 16.03 million 150 bp reads were mapped for each sample (Figure 2A). Due to the random priming process of lcWGS, a more diverse set of DNA fragments were sequenced. This enabled lcWGS sequences to cover a wider range of the genome at 18.28% per sample on average, but with a lower mean coverage of 1.39x at each capture site (Figure 2B, 2C). This resulted in a mean mapped coverage of 0.27x per sample genome-wide (Figure 2D).

### Genotype Statistics

In our genotyping pipeline, we imputed a total of 10,684,883 bi-allelic SNPs. Following the quality control procedures outlined in the methods section, 7,323,260 SNPs were retained. Out of these retained SNPs, 7,148,654 were located on autosomal chromosomes, 174,374 were on chromosome X, 118 were on chromosome Y, and 114 were on mitochondria (Figure 3).

**Figure 3.**
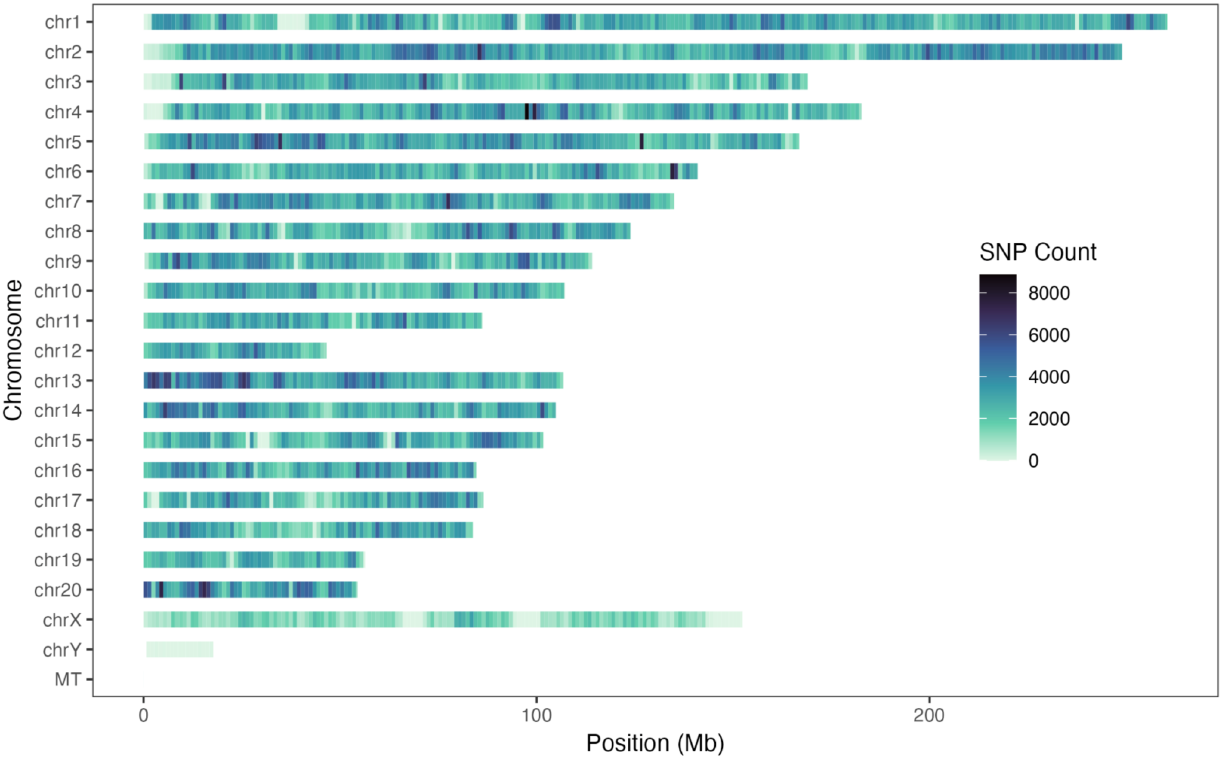
Imputed bi-allelic SNPs density distribution heatmap on each chromosome with one-megabase windows.

Among the 7,148,654 SNPs on autosomes, 1,602,374 were found to be monomorphic with a minor allele frequency (**MAF**) of 0. We assume that these SNPs, which were polymorphic in the HS founders, became monomorphic in the outbred HS population due to genetic drift (Munro et al. 2022). 183,621 were rare SNPs with MAF ≤ 0.005, which means there were 5,362,659 common SNPs with MAF > 0.005 (Figure S5A in Supplementary Material). Out of the 7,148,654 SNPs, 39,606 violated Hardy-Weinberg Equilibrium (**HWE**) with a -log10(p-value) ≥ 10 (arbitrary cutoff, Figure S5B in Supplementary Material), and 36,664 had a genotype missing rate higher than 0.1 (Figure S5C in Supplementary Material). Consequently, a total of 5,292,916 autosomal SNPs had a MAF > 0.005, HWE -log10 p-value < 10 and missing rate ≤ 0.1.

### Sex Chromosomes

Due to the different inheritance patterns on sex chromosomes in males and females, we investigated the SNPs on chromosome X and Y separately in each sex. Among the 7,222 female samples included in this study, we observed that out of the 174,374 SNPs on chromosome X, 47,882 were monomorphic, 1,375 were rare, and 125,117 were common (Figure S6A in Supplementary Material). 627 SNPs violated HWE (Figure S6B in Supplementary Material), and 582 SNPs had a missing rate higher than 0.1 (Figure S6C in Supplementary Material). This led to a total of 123,997 chromosome X SNPs for females with a MAF > 0.005, HWE -log10 p-value < 10 and missing rate ≤ 0.1. Chromosome Y SNPs were discarded for female samples. In the 7,283 male samples used in this study, among the 174,374 SNPs on chromosome X, 46,319 were monomorphic, 3,227 were rare, and 124,828 were common (Figure S6D in Supplementary Material). Because males have only one copy of the X chromosome, we did not test them for HWE, but we found 2,223 chromosome X SNPs had a missing rate higher than 0.1 (Figure S6E in Supplementary Material). This resulted in a total of 122,693 chromosome X SNPs for males with a MAF > 0.005 and missing rate ≤ 0.1. The 118 SNPs on chromosome Y for male samples had a missing rate ≤ 0.1, but they were all monomorphic SNPs with a MAF of 0.

Out of the 114 SNPs on the mitochondrial chromosome, 30 were found to be monomorphic with a MAF of 0, and the remaining 74 were common SNPs with MAF > 0.005 (Figure S7A in Supplementary Material). HWE was also not tested for mitochondrial SNPs, but all of them had a genotype missing rate lower than 0.1 (Figure S7B in Supplementary Material). Consequently, a total of 74 mitochondria SNPs had a MAF > 0.005 and missing rate ≤ 0.1.

We have recently published a separate paper that uses the same genotypes described here to examine Y and mitochondrial chromosome haplogroups (Okamoto et al. 2023).

### Genotype Accuracy

As described in the methods section, in the 15,552 outbred HS rats we genotyped, there were 88 outbred HS rats that had been sequenced with ddGBS, lcWGS and 33.26x high coverage WGS. We tested our genotyping pipeline’s accuracy by comparing genotypes imputed from ddGBS and lcWGS with SNPs called using high coverage WGS without any imputation, which we refer to as the ‘truth set’. Specifically, for each sample, we looked at the concordance rate of overlap and non-missing SNPs between the imputed genotypes and the truth set. On average, 5,429,453 polymorphic SNPs were shared between imputation from ddGBS sequences and variant calling from 33.26x high coverage WGS, with a mean concordance rate of 99.76% (Figure 4). Similarly, we observed that 5.43 million SNPs were shared between lcWGS and 33.26x high coverage WGS, with a mean concordance rate of 99.78% (Figure 4).

**Figure 4.**
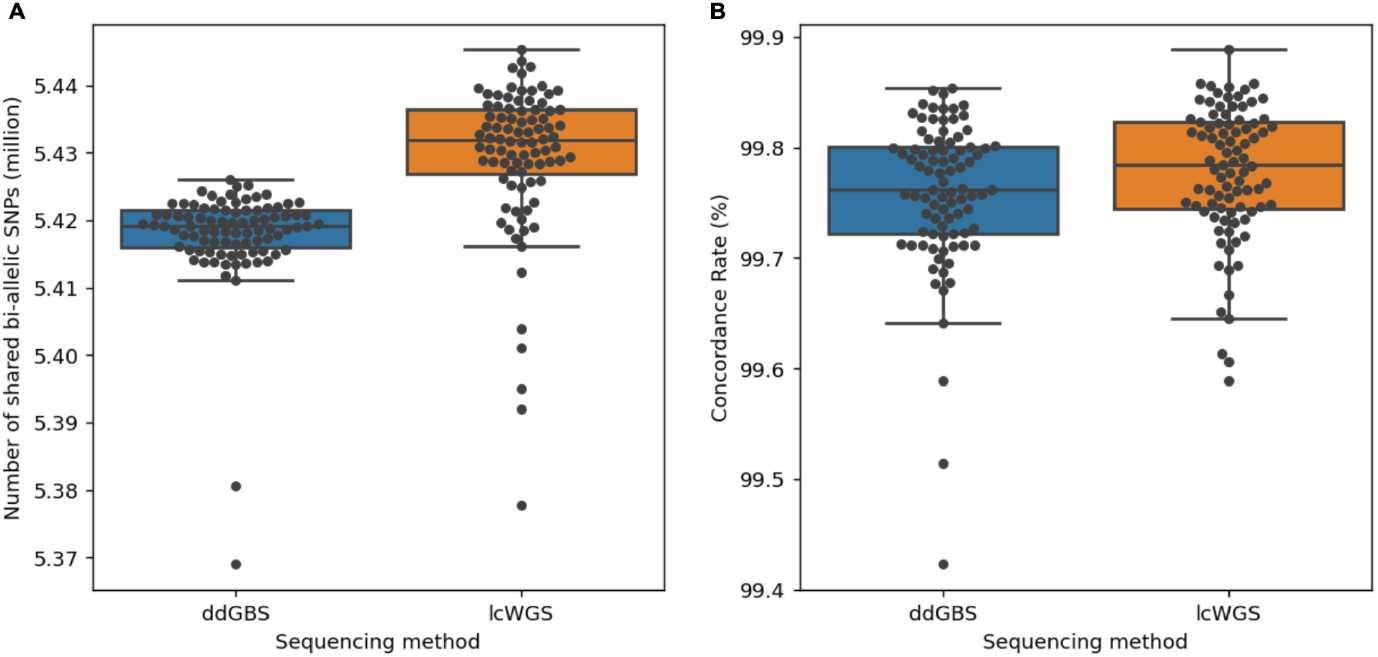
Imputed genotypes demonstrate high concordance with 33.26x high coverage WGS for millions of bi-allelic SNPs. **A**. Number of bi-allelic SNPs compared (million). ddGBS mean: 5.42, SD: 0.01; lcWGS mean: 5.43, SD: 0.01. **B**. Concordance rate with 33.26x high coverage WGS (%). ddGBS mean: 99.76, SD: 0.07; lcWGS mean: 99.78, SD: 0.06.

### Batch Effects on ddGBS and lcWGS Genotypes

To investigate potential batch effects of different sequencing methods, we performed a principal component analysis on the autosomal genotypes of the 88 HS outbred rats sequenced with both ddGBS and lcWGS (Figure 5). Overlapping first and second principal component (PC) values without apparent clustering between the two methods indicate that both methods capture equivalent information from the genome, meaning there are no obvious method-specific batch effects introduced by the pipeline. Additionally, we did not observe any batch effects in any other PCs that explained more than 10% of the variance (Figure S8 in Supplementary Material).

**Figure 5.**
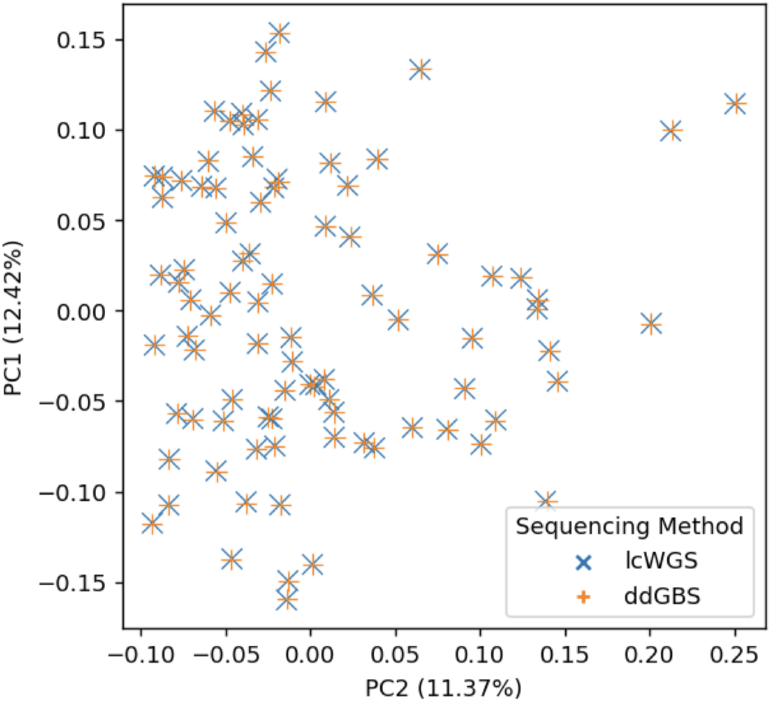
Overlapping first and second principal components on genotypes shows no batch effects between different sequencing methods.

## Discussion

While large-scale genetic studies in humans often use genotyping microarrays and imputation, similar resources are not available for most other species. Although there are examples where human genetic studies use low coverage WGS and imputation for genotyping, they typically require higher coverage because of more diverse and smaller haplotype blocks (Cai et al. 2015; Petter et al. 2020; Li et al. 2021; Wasik et al. 2021; Li et al. 2024). Our genotyping method takes advantage of the unique HS population structure caused by interbreeding eight inbred founders. Because the founders are fully sequenced, we are able to construct a high quality reference panel for HS rats, which enables us to achieve highly accurate imputation for their genotypes even with low read coverage (0.21x mean ddGBS and 0.27x mean lcWGS).

Others have reported a similar genotyping strategy of using GBS or lcWGS alone and imputation for AIL and CFW mice (Nicod et al. 2016; Parker et al. 2016; Gonzales et al. 2018). Nicod et al. used lcWGS sequence data with STITCH imputation on CFW mice. Parker et al. used GBS sequence data with IMPUTE2 on CFW mice. Gonzales et al. used GBS sequence data with BEAGLE on AIL mice. Their estimated genotype concordance rates were 98.1%, 97.0% and 96.96% respectively compared to the MegaMUGA array. Our previous work of using ddGBS and imputation (two rounds of imputation: BEAGLE and IMPUTE2) to genotype HS rats was able to produce over 3.7 million SNPs with a concordance rate of 99.0% compared to a custom Affymetrix Axiom MiRat 625k microarray (Gileta et al. 2020). These four studies also included a variant calling step to identify candidate variants using either ANGSD or GATK before imputation. Our genotyping method described here doesn’t require such a variant calling step, which reduces computation. Our method combines ddGBS and lcWGS sequence data and uses STITCH in conjunction with a fully sequenced founder reference panel to achieve genotype imputation. As a result, we achieve a high genotype concordance rate (>99.76%) compared to high-coverage (33.26x coverage) WGS.

Our genotyping method provides a robust method for genotyping HS rats by effectively imputing SNP genotypes from two different sequencing protocols without significant batch effects. Even with low read coverage, our method produced highly accurate genotypes. Our method is cost-effective due to the previously developed affordable ddGBS technique and low-cost lcWGS, which use commercially available library preparation kits and liquid handling robots, improving throughput. Additionally, our method combines ddGBS and lcWGS sequences for genotype imputation, enabling old ddGBS genotyped rats to be analyzed in tandem with more recently genotyped HS rats.

The differences we observed in aligned sequence statistics between ddGBS and lcWGS (Figure 2) reflect the different nature of the DNA sequences captured by two sequencing methods. Double restriction enzyme digestion limits ddGBS to only capture the DNA fragments near the enzyme cut sites, while random priming helps lcWGS capture DNA fragments across the genome randomly. Despite the differences in captured DNA fragments, the genotype concordances of imputed SNPs for both ddGBS and lcWGS are remarkably high, at 99.76% and 99.78%, respectively. This concordance demonstrates the strength of our pipeline in producing high-accuracy genotypes in HS rats, which provides a strong foundation for genetic studies in this population.

The GBS sequencing method was originally developed by Elshire et al. (Elshire et al. 2011) and modified to accommodate other species such as soybean (Sonah et al. 2013), rice (Furuta et al. 2017), oat (Fu 2018), chicken (Pértille et al. 2016; Wang et al. 2017), fox (Johnson et al. 2015), and cattle (Donato et al. 2013), and mouse (Parker et al. 2016; Gonzales et al. 2018). Our lab modified GBS for use in HS rats (Gileta et al. 2020). In this study, we further improved our genotyping methods by harmonizing the previously produced ddGBS sequences and newly sequenced lcWGS sequences with commercial WGS technique in support of large scale genetic studies. The principals of our genotyping method can be easily adapted for use in other populations, especially for those in which the founders are fully sequenced.

In summary, we developed a genotyping method for HS rats that is both cost-effective and high-throughput, yielding highly accurate genotypes. Our method can be readily applied to other species with minimal adjustments, forming a basis for conducting extensive genetic research in non-human species.

## Data Availability

HS rats are available at https://ratgenes.org/cores/core-b/. Wet lab procedures are documented in protocols.io https://www.protocols.io/workspaces/cgord (spleen cutting: http://dx.doi.org/10.17504/protocols.io.36wgq7nryvk5/v1, DNA extraction: http://dx.doi.org/10.17504/protocols.io.8epv59reng1b/v1, normalization and randomization: http://dx.doi.org/10.17504/protocols.io.261genw5dg47/v1, library preparation: http://dx.doi.org/10.17504/protocols.io.j8nlkkm85l5r/v1, pooling and sequencing: http://dx.doi.org/10.17504/protocols.io.yxmvmnw29g3p/v1). Raw sequencing reads for ddGBS and lcWGS are available in NCBI SRA: PRJNA1022514. Eight HS inbred founders WGS raw reads are available in NCBI SRA: PRJNA487943 and PRJNA1048943. 88 selected HS rats WGS raw reads are available in NCBI SRA: PRJNA107614. High coverage WGS GATK genotyping pipeline code is available in Zenodo https://doi.org/10.5281/zenodo.6584834 and Github https://github.com/Palmer-Lab-UCSD/High-Coverage-WGS-GATK-Genotyping-Pipeline. High coverage WGS DeepVariant genotyping pipeline code is available in Zenodo https://doi.org/10.5281/zenodo.10027133 and Github https://github.com/Palmer-Lab-UCSD/High-Coverage-WGS-DeepVariant-Genotyping-Pipeline. Genotyping pipeline and analysis code is available in Zenodo https://doi.org/10.5281/zenodo.10002191 and GitHub https://github.com/Palmer-Lab-UCSD/HS-Rats-Genotyping-Pipeline. Genotype data after quality control are available in UC San Diego Library Digital Collections https://doi.org/10.6075/J0445MPC.

## Acknowledgments

This publication includes data generated at the UC San Diego IGM Genomics Center utilizing an Illumina NovaSeq 6000 that was purchased with funding from a National Institutes of Health SIG grant (#S10 OD026929) San Diego Supercomputer Center (2022): Triton Shared Computing Cluster. University of California, San Diego. Service. https://doi.org/10.57873/T34W2R

## References

Abdellaoui A, Yengo L, Verweij KJH, Visscher PM. 2023. 15 years of GWAS discovery: Realizing the promise. Am J Hum Genet. 110(2):179–194. doi:10.1016/j.ajhg.2022.12.011.

Aganezov S, Yan SM, Soto DC, Kirsche M, Zarate S, Avdeyev P, Taylor DJ, Shafin K, Shumate A, Xiao C, et al. 2022. A complete reference genome improves analysis of human genetic variation. Science. 376(6588):eabl3533. doi:10.1126/science.abl3533.

Alliance of Genome Resources Consortium. 2022. Harmonizing model organism data in the Alliance of Genome Resources. Genetics. 220(4):iyac022. doi:10.1093/genetics/iyac022.

Baud A, Hermsen R, Guryev V, Stridh P, Graham D, McBride MW, Foroud T, Calderari S, Diez M, Ockinger J, et al. 2013. Combined sequence-based and genetic mapping analysis of complex traits in outbred rats. Nat Genet. 45(7):767–775. doi:10.1038/ng.2644.

Broad Institute. 2019. Picard Toolkit. [accessed 2023 Apr 10]. https://broadinstitute.github.io/picard/.

Bushnell B. BBTools. http://sourceforge.net/projects/bbmap/.

Cai N, Bigdeli TB, Kretzschmar W, Li Y, Liang J, Song L, Hu J, Li Q, Jin W, Hu Z, et al. 2015. Sparse whole-genome sequencing identifies two loci for major depressive disorder. Nature. 523(7562):588–591. doi:10.1038/nature14659.

Chitre AS, Hebda-Bauer EK, Blandino P, Bimschleger H, Nguyen K-M, Maras P, Li F, Ozel AB, Pan Y, Polesskaya O, et al. 2023. Genome-wide association study in a rat model of temperament identifies multiple loci for exploratory locomotion and anxiety-like traits. Front Genet. 13. doi:10.3389/fgene.2022.1003074. [accessed 2024 May 22].

https://www.frontiersin.org/journals/genetics/articles/10.3389/fgene.2022.1003074/full.

Chitre AS, Polesskaya O, Holl K, Gao J, Cheng R, Bimschleger H, Garcia Martinez A, George T, Gileta AF, Han W, et al. 2020. Genome-Wide Association Study in 3,173 Outbred Rats Identifies Multiple Loci for Body Weight, Adiposity, and Fasting Glucose. Obesity. 28(10):1964–1973. doi:10.1002/oby.22927.

Danecek P, Bonfield JK, Liddle J, Marshall J, Ohan V, Pollard MO, Whitwham A, Keane T, McCarthy SA, Davies RM, et al. 2021. Twelve years of SAMtools and BCFtools. GigaScience. 10(2):giab008. doi:10.1093/gigascience/giab008.

Davies RW, Flint J, Myers S, Mott R. 2016. Rapid genotype imputation from sequence without reference panels. Nat Genet. 48(8):965–969. doi:10.1038/ng.3594.

Didion JP, Yang H, Sheppard K, Fu C-P, McMillan L, de Villena FP-M, Churchill GA. 2012. Discovery of novel variants in genotyping arrays improves genotype retention and reduces ascertainment bias. BMC Genomics. 13(1):34. doi:10.1186/1471-2164-13-34.

Donato MD, Peters SO, Mitchell SE, Hussain T, Imumorin IG. 2013. Genotyping-by-Sequencing (GBS): A Novel, Efficient and Cost-Effective Genotyping Method for Cattle Using Next-Generation Sequencing. PLOS ONE. 8(5):e62137. doi:10.1371/journal.pone.0062137.

Elshire RJ, Glaubitz JC, Sun Q, Poland JA, Kawamoto K, Buckler ES, Mitchell SE. 2011. A Robust, Simple Genotyping-by-Sequencing (GBS) Approach for High Diversity Species. Orban L, editor. PLoS ONE. 6(5):e19379. doi:10.1371/journal.pone.0019379.

Fowler S, Wang T, Munro D, Kumar A, Chitre AS, Hollingsworth TJ, Garcia Martinez A, St. Pierre CL, Bimschleger H, Gao J, et al. 2023. Genome-wide association study finds multiple loci associated with intraocular pressure in HS rats. Front Genet. 13. doi:10.3389/fgene.2022.1029058. [accessed 2024 May 22]. https://www.frontiersin.org/journals/genetics/articles/10.3389/fgene.2022.1029058/full.

Frazer KA, Ballinger DG, Cox DR, Hinds DA, Stuve LL, Gibbs RA, Belmont JW, Boudreau A, Hardenbol P, Leal SM, et al. 2007. A second generation human haplotype map of over 3.1 million SNPs. Nature. 449(7164):851–861. doi:10.1038/nature06258.

Fu Y-B. 2018. Oat evolution revealed in the maternal lineages of 25 Avena species. Sci Rep. 8(1):4252. doi:10.1038/s41598-018-22478-4.

Furuta T, Ashikari M, Jena KK, Doi K, Reuscher S. 2017. Adapting Genotyping-by-Sequencing for Rice F2 Populations. G3 GenesGenomesGenetics. 7(3):881–893. doi:10.1534/g3.116.038190.

Gileta AF, Fitzpatrick CJ, Chitre AS, Pierre CLS, Joyce EV, Maguire RJ, McLeod AM, Gonzales NM, Williams AE, Morrow JD, et al. 2022. Genetic characterization of outbred Sprague Dawley rats and utility for genome-wide association studies. PLOS Genet. 18(5):e1010234. doi:10.1371/journal.pgen.1010234.

Gileta AF, Gao J, Chitre AS, Bimschleger HV, St. Pierre CL, Gopalakrishnan S, Palmer AA. 2020. Adapting Genotyping-by-Sequencing and Variant Calling for Heterogeneous Stock Rats. G3 GenesGenomesGenetics. 10(7):2195–2205. doi:10.1534/g3.120.401325.

Gonzales NM, Seo J, Hernandez Cordero AI, St. Pierre CL, Gregory JS, Distler MG, Abney M, Canzar S, Lionikas A, Palmer AA. 2018. Genome wide association analysis in a mouse advanced intercross line. Nat Commun. 9(1):5162. doi:10.1038/s41467-018-07642-8.

Gunturkun MH, Wang T, Chitre AS, Garcia Martinez A, Holl K, St. Pierre C, Bimschleger H, Gao J, Cheng R, Polesskaya O, et al. 2022. Genome-Wide Association Study on Three Behaviors Tested in an Open Field in Heterogeneous Stock Rats Identifies Multiple Loci Implicated in Psychiatric Disorders. Front Psychiatry. 13. doi:10.3389/fpsyt.2022.790566. [accessed 2024 May 22]. https://www.frontiersin.org/journals/psychiatry/articles/10.3389/fpsyt.2022.790566/full.

Hannon Lab. 2010. FASTX-Toolkit. [accessed 2023 Apr 10]. http://hannonlab.cshl.edu/fastx_toolkit/.

Johannesson M, Lopez-Aumatell R, Stridh P, Diez M, Tuncel J, Blázquez G, Martinez-Membrives E, Cañete T, Vicens-Costa E, Graham D, et al. 2009. A resource for the simultaneous high-resolution mapping of multiple quantitative trait loci in rats: The NIH heterogeneous stock. Genome Res. 19(1):150–158. doi:10.1101/gr.081497.108.

Johnson JL, Wittgenstein H, Mitchell SE, Hyma KE, Temnykh SV, Kharlamova AV, Gulevich RG, Vladimirova AV, Fong HWF, Acland GM, et al. 2015. Genotyping-By-Sequencing (GBS) Detects Genetic Structure and Confirms Behavioral QTL in Tame and Aggressive Foxes (Vulpes vulpes). PLOS ONE. 10(6):e0127013. doi:10.1371/journal.pone.0127013.

Li H. 2013. Aligning sequence reads, clone sequences and assembly contigs with BWA-MEM. [accessed 2023 Apr 10]. http://arxiv.org/abs/1303.3997.

Li JH, Findley K, Pickrell JK, Blease K, Zhao J, Kruglyak S. 2024. Low-pass sequencing plus imputation using avidity sequencing displays comparable imputation accuracy to sequencing by synthesis while reducing duplicates. G3 GenesGenomesGenetics. 14(2):jkad276. doi:10.1093/g3journal/jkad276.

Li JH, Mazur CA, Berisa T, Pickrell JK. 2021. Low-pass sequencing increases the power of GWAS and decreases measurement error of polygenic risk scores compared to genotyping arrays. Genome Res. 31(4):529–537. doi:10.1101/gr.266486.120.

Marchini J, Howie B. 2010. Genotype imputation for genome-wide association studies. Nat Rev Genet. 11(7):499–511. doi:10.1038/nrg2796.

Martin M. 2011. Cutadapt removes adapter sequences from high-throughput sequencing reads. EMBnet.journal. 17(1):10–12. doi:10.14806/ej.17.1.200.

McVean GA, Altshuler (Co-Chair) DM, Durbin (Co-Chair) RM, Abecasis GR, Bentley DR, Chakravarti A, Clark AG, Donnelly P, Eichler EE, Flicek P, et al. 2012. An integrated map of genetic variation from 1,092 human genomes. Nature. 491(7422):56–65. doi:10.1038/nature11632.

Munro D, Wang T, Chitre AS, Polesskaya O, Ehsan N, Gao J, Gusev A, Woods LCS, Saba LM, Chen H, et al. 2022. The regulatory landscape of multiple brain regions in outbred heterogeneous stock rats. Nucleic Acids Res. 50(19):10882–10895. doi:10.1093/nar/gkac912.

Nicod J, Davies RW, Cai N, Hassett C, Goodstadt L, Cosgrove C, Yee BK, Lionikaite V, McIntyre RE, Remme CA, et al. 2016. Genome-wide association of multiple complex traits in outbred mice by ultra-low-coverage sequencing. Nat Genet. 48(8):912–918. doi:10.1038/ng.3595.

Okamoto F, Chitre AS, Missfeldt Sanches T, Chen D, Munro D, Polesskaya O, Palmer AA. 2023 Nov 29. Y and Mitochondrial Chromosomes in the Heterogeneous Stock Rat Population. bioRxiv.:2023.11.29.566473. doi:10.1101/2023.11.29.566473.

Palmer RHC, Johnson EC, Won H, Polimanti R, Kapoor M, Chitre A, Bogue MA, Benca-Bachman CE, Parker CC, Verma A, et al. 2021. Integration of evidence across human and model organism studies: A meeting report. Genes Brain Behav. 20(6):e12738. doi:10.1111/gbb.12738.

Parker CC, Gopalakrishnan S, Carbonetto P, Gonzales NM, Leung E, Park YJ, Aryee E, Davis J, Blizard DA, Ackert-Bicknell CL, et al. 2016. Genome-wide association study of behavioral, physiological and gene expression traits in outbred CFW mice. Nat Genet. 48(8):919–926. doi:10.1038/ng.3609.

Parker CC, Philip VM, Gatti DM, Kasparek S, Kreuzman AM, Kuffler L, Mansky B, Masneuf S, Sharif K, Sluys E, et al. 2022. Genome-wide association mapping of ethanol sensitivity in the Diversity Outbred mouse population. Alcohol Clin Exp Res. 46(6):941–960. doi:10.1111/acer.14825.

Pértille F, Guerrero-Bosagna C, Silva VH da, Boschiero C, Nunes J de R da S, Ledur MC, Jensen P, Coutinho LL. 2016. High-throughput and Cost-effective Chicken Genotyping Using Next-Generation Sequencing. Sci Rep. 6(1):26929. doi:10.1038/srep26929.

Petter E, Schweiger R, Shahino B, Shor T, Aker M, Almog L, Weissglas-Volkov D, Naveh Y, Navon O, Carmi S, et al. 2020. Relative matching using low coverage sequencing. :2020.09.09.289322. doi:10.1101/2020.09.09.289322. [accessed 2023 Sep 19]. https://www.biorxiv.org/content/10.1101/2020.09.09.289322v3.

Ramdas S, Ozel AB, Treutelaar MK, Holl K, Mandel M, Woods LCS, Li JZ. 2019. Extended regions of suspected mis-assembly in the rat reference genome. Sci Data. 6(1):39. doi:10.1038/s41597-019-0041-6.

Solberg Woods LC, Palmer AA. 2019. Using Heterogeneous Stocks for Fine-Mapping Genetically Complex Traits. In: Hayman GT, Smith JR, Dwinell MR, Shimoyama M, editors. Rat Genomics. New York, NY: Springer. p. 233–247. [accessed 2024 May 22]. 10.1007/978-1-4939-9581-3_11.

Sonah H, Bastien M, Iquira E, Tardivel A, Légaré G, Boyle B, Normandeau É, Laroche J, Larose S, Jean M, et al. 2013. An Improved Genotyping by Sequencing (GBS) Approach Offering Increased Versatility and Efficiency of SNP Discovery and Genotyping. PLOS ONE. 8(1):e54603. doi:10.1371/journal.pone.0054603.

Tim F, Nils H. 2023. fgbio. [accessed 2023 Apr 10]. https://github.com/fulcrumgenomics/fgbio.

Uffelmann E, Huang QQ, Munung NS, de Vries J, Okada Y, Martin AR, Martin HC, Lappalainen T, Posthuma D. 2021. Genome-wide association studies. Nat Rev Methods Primer. 1(1):1–21. doi:10.1038/s43586-021-00056-9.

Van der Auwera GA, O’Connor BD. 2020. Genomics in the Cloud: Using Docker, GATK, and WDL in Terra. 1st ed. O’Reilly Media.

Wang Y, Cao X, Zhao Y, Fei J, Hu X, Li N. 2017. Optimized double-digest genotyping by sequencing (ddGBS) method with high-density SNP markers and high genotyping accuracy for chickens. PLOS ONE. 12(6):e0179073. doi:10.1371/journal.pone.0179073.

Wasik K, Berisa T, Pickrell JK, Li JH, Fraser DJ, King K, Cox C. 2021. Comparing low-pass sequencing and genotyping for trait mapping in pharmacogenetics. BMC Genomics. 22(1):197. doi:10.1186/s12864-021-07508-2.

Welter D, MacArthur J, Morales J, Burdett T, Hall P, Junkins H, Klemm A, Flicek P, Manolio T, Hindorff L, et al. 2014. The NHGRI GWAS Catalog, a curated resource of SNP-trait associations. Nucleic Acids Res. 42(D1):D1001–D1006. doi:10.1093/nar/gkt1229.

Woods LCS, Mott R. 2017. Heterogeneous Stock Populations for Analysis of Complex Traits. In: Schughart K, Williams RW, editors. Systems Genetics: Methods and Protocols. New York, NY: Springer. (Methods in Molecular Biology). p. 31–44. [accessed 2023 Apr 6]. 10.1007/978-1-4939-6427-7_2.

Zhou X, St. Pierre CL, Gonzales NM, Zou J, Cheng R, Chitre AS, Sokoloff G, Palmer AA. 2020. Genome-Wide Association Study in Two Cohorts from a Multi-generational Mouse Advanced Intercross Line Highlights the Difficulty of Replication Due to Study-Specific Heterogeneity. G3 GenesGenomesGenetics. 10(3):951–965. doi:10.1534/g3.119.400763.

Zou J, Gopalakrishnan S, Parker CC, Nicod J, Mott R, Cai N, Lionikas A, Davies RW, Palmer AA, Flint J. 2022. Analysis of independent cohorts of outbred CFW mice reveals novel loci for behavioral and physiological traits and identifies factors determining reproducibility. G3 GenesGenomesGenetics. 12(1):jkab394. doi:10.1093/g3journal/jkab394.

