## Supplementary Material for "A Cost-effective, High-throughput, Highly Accurate Genotyping Method for Outbred Populations"

**Supplementary Method S1.** High coverage WGS sequencing

The Agencourt DNAdvance Kit (Beckman Coulter Life Sciences, Indianapolis, IN) was used to extract DNA from the small intestine or liver tissues of eight inbred HS rat founders, and from the spleen tissues of 88 outbred HS rats. Next, DNA libraries were prepared according to the manufacturer's protocol using KAPA HyperPrep Kits (Roche, Basel, Switzerland). The DNA were then sequenced with Illumina NovaSeq 6000 to obtain 33.26x coverage 150 bp paired-end reads at the University of California San Diego Institute for Genomic Medicine Genomics Center.

**Supplementary Method S2.** High coverage WGS GATK genotyping

SNPs and indels from the eight inbred rat founder strains and 88 outbred HS rats were jointly called using the Base Quality Score Recalibration bootstrapping strategy with Genome Analysis Toolkit (**GATK**) HaplotypeCaller and GenotypeGVCFs as shown in Supplementary Figure S1 (Van der Auwera and O'Connor 2020). At the sequence quality control step, BBDuk was used to trim the adapters and poly-G tails with length greater than 50 bp (Bushnell). We then aligned the sequences to the reference genome and did quality control on aligned reads with BWA-mem and Samtools (Li 2013; Danecek et al. 2021). The called SNPs were filtered out with Qual < 30, QualByDepth < 5, FisherStrand > 60, StrandOddsRatio > 3, RMSMappingQuality < 40, MappingQualityRankSumTest < -12.5, ReadPosRankSumTest < -8, and the indels with Qual < 30, QualByDepth < 5, FisherStrand > 200, ReadPosRankSumTest < -20. The source code of the high coverage WGS GATK genotyping pipeline can be found in the Palmer Lab GitHub repository (https://github.com/Palmer-Lab-UCSD/High-Coverage-WGS-Genotyping-Pipeline, DOI: https://doi.org/10.5281/zenodo.6584834).

**Supplementary Figure S1**. High coverage WGS GATK genotyping pipeline flowchart.

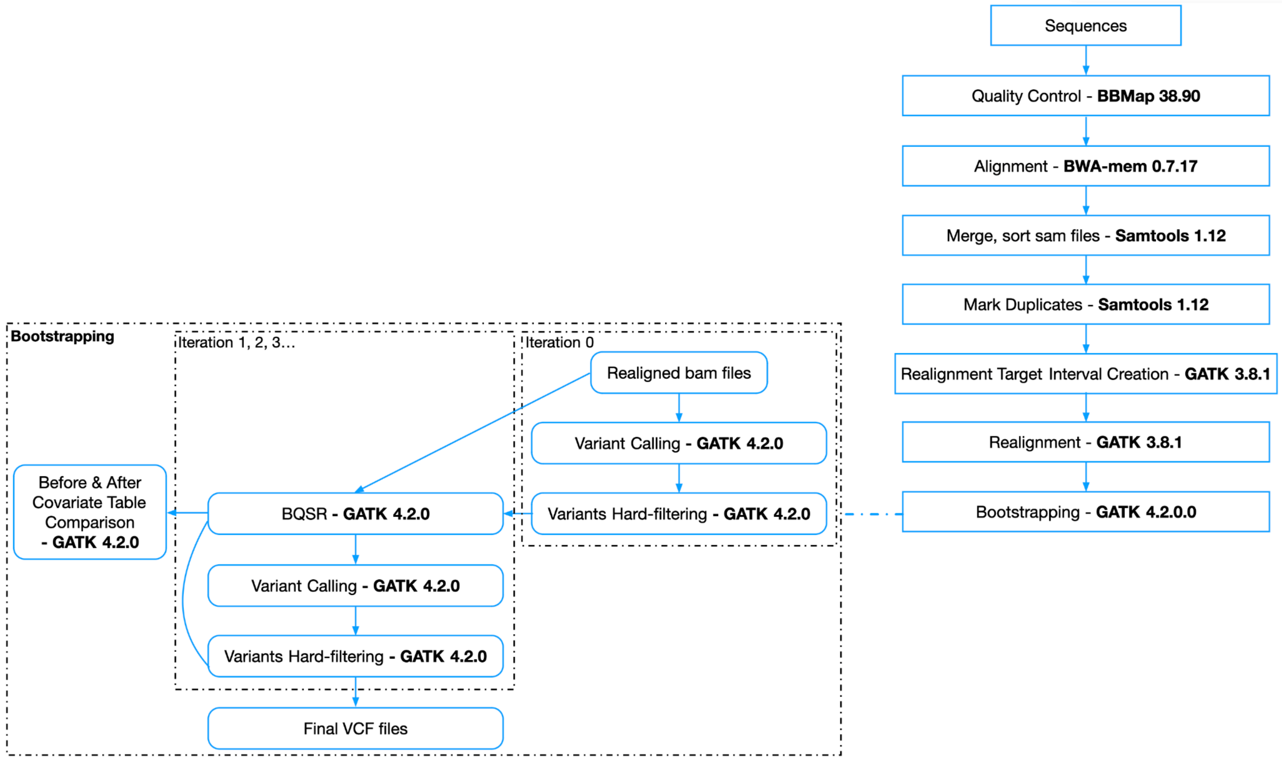

**Supplementary Method S3.** High coverage WGS DeepVariant genotyping pipeline

SNPs and indels from the eight inbred rat founder strains were jointly called using DeepVariant and GLnexus based on the aligned reads after realignment from high coverage WGS GATK genotyping process as shown in Supplementary Figure S2 (Poplin et al. 2018; Yun et al. 2021). The source code of the high coverage WGS GATK genotyping pipeline can be found in the Palmer Lab GitHub repository (https://github.com/Palmer-Lab-UCSD/High-Coverage-WGS-DeepVariant-Genotyping-Pipeline, DOI: https://doi.org/10.5281/zenodo.10027133).

**Supplementary Figure S2.** High coverage WGS DeepVariant genotyping pipeline flowchart.

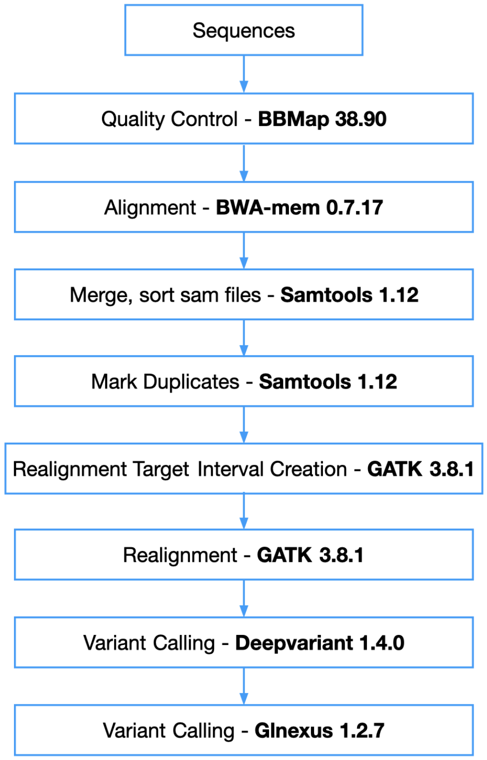

**Supplementary Figure S3**. Ratio of mapped reads on chromosome X and Y. Green line: y=x/20.5+0.07, red lines: y=x/20.5+0.07±0.02. Females above the top red line and males below the bottom red line are marked as failures on sex quality control. Samples between two read lines are marked as suspect on sex quality control. All suspect samples also failed other quality control procedures and were excluded in the final set of samples. **A.** ddGBS and lcWGS samples (7,797 males: 7,745 pass, 13 suspect, 39 fail; 7,755 females: 7,689 pass, 19 suspect, 47 fail). **B.** ddGBS samples (3,219 males: 3,209 pass, 0 suspect, 10 fail; 3,160 females: 3,150 pass, 0 suspect, 10 fail). **C.** lcWGS samples (4,578 males: 4,536 pass, 13 suspect, 29 fail; 4,595 females: 4,539 pass, 19 suspect, 37 fail).

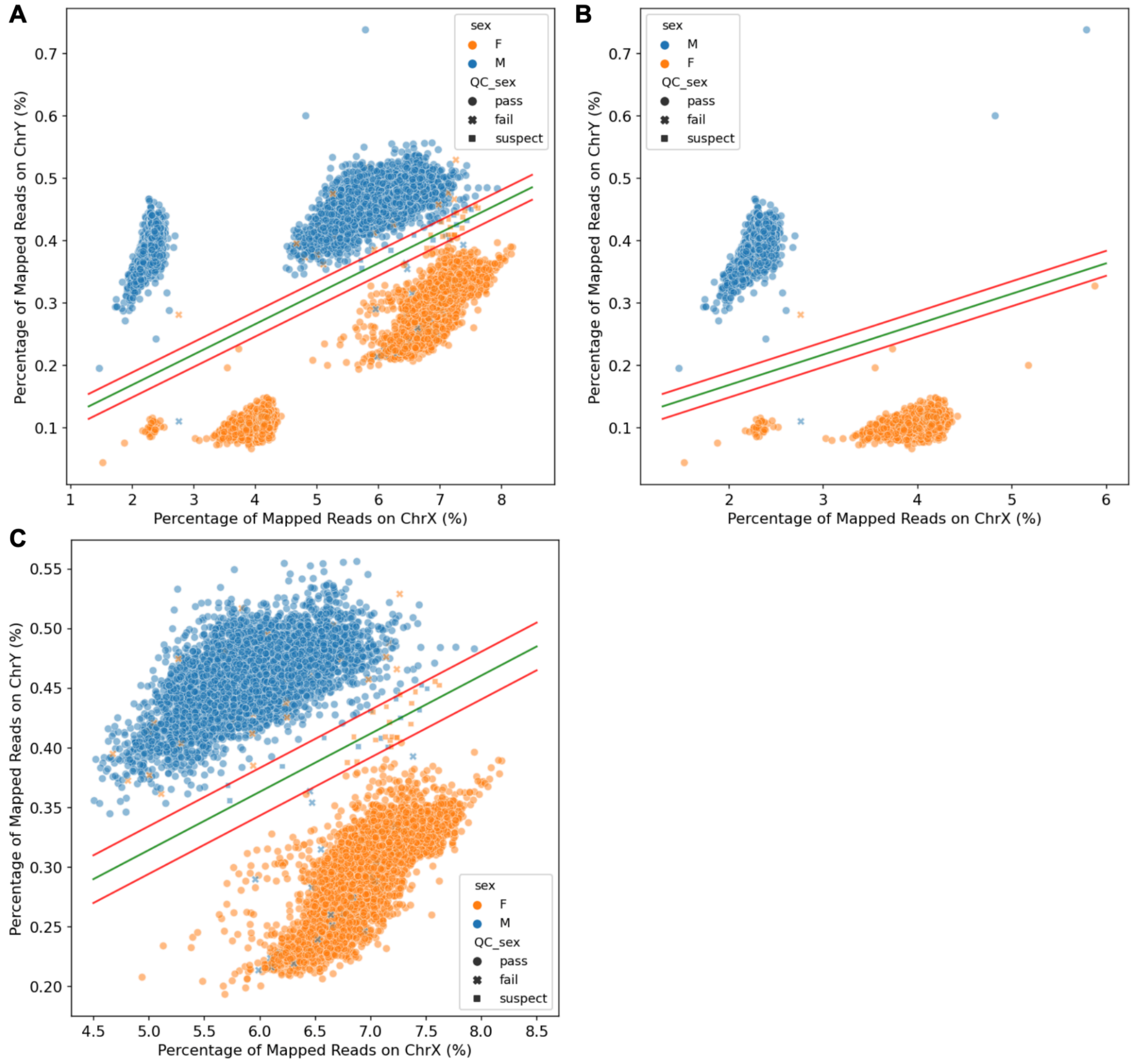

**Supplementary Figure S4**. Sample genotype heterozygosity rate vs. sample genotype missing rate. Yellow line indicates the mean heterozygosity rate + four standard deviations, orange line indicates the mean heterozygosity rate - four standard deviations, red line indicates sample genotype missing rate 0.1. Samples with a genotype missing rate exceeding 0.1 or a genotype heterozygosity rate falling outside the range of the mean ± four standard deviations were marked as failures. **A.** ddGBS samples (6,379 samples: 6,345 pass, 34 fail). **B.** lcWGS samples (9,173 samples: 9,054 pass, 119 fail).

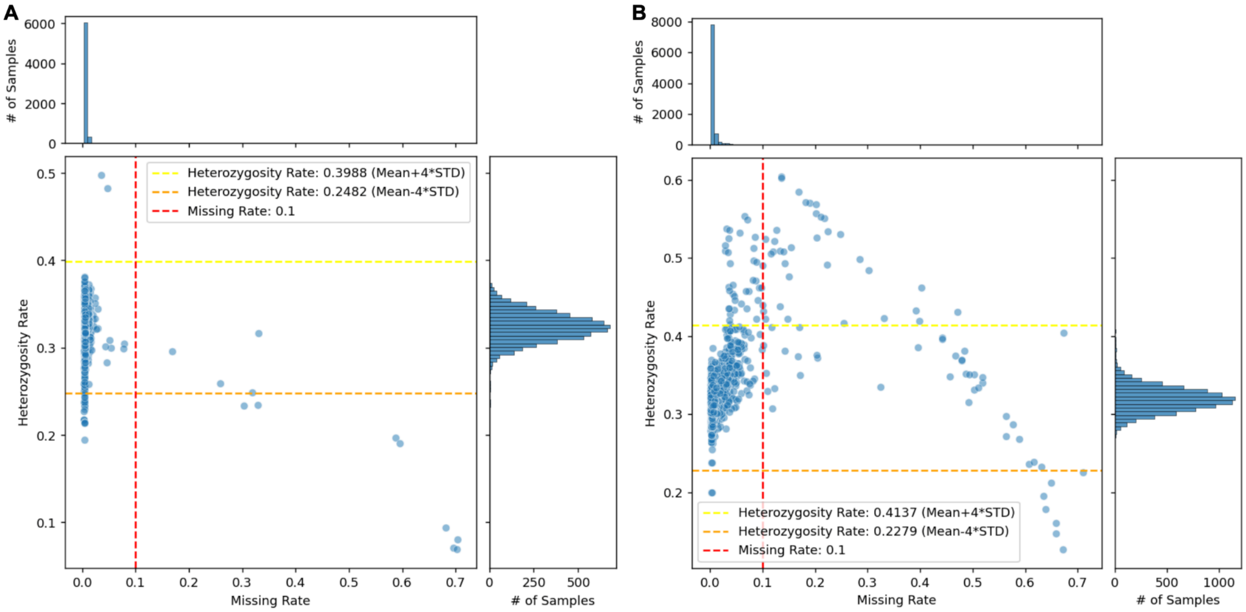

**Supplementary Figure S5.** Autosomal SNPs statistics. **A.** Autosomal SNPs minor allele frequency histogram. **B.** Autosomal SNPs Hardy-Weinberg Equilibrium -log10 p-value histogram. **C.** Autosomal SNPs missing rate histogram.

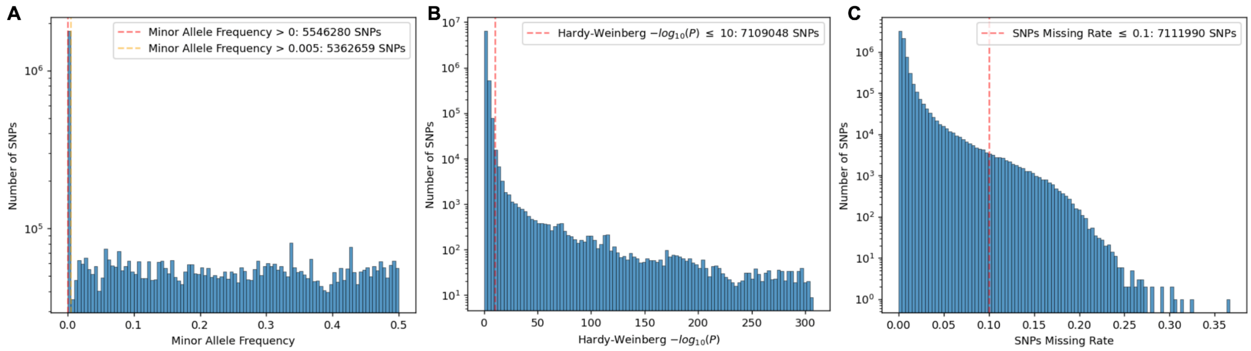


**Supplementary Figure S6.** Chromosome X SNPs statistics. **A.** Chromosome X SNPs minor allele frequency histogram in female samples. **B.** Chromosome X SNPs Hardy-Weinberg Equilibrium -log10 p-value histogram in female samples. **C.** Chromosome X SNPs missing rate histogram in female samples. **D.** Chromosome X SNPs minor allele frequency histogram in male samples. **E.** Chromosome X SNPs missing rate histogram in male samples.

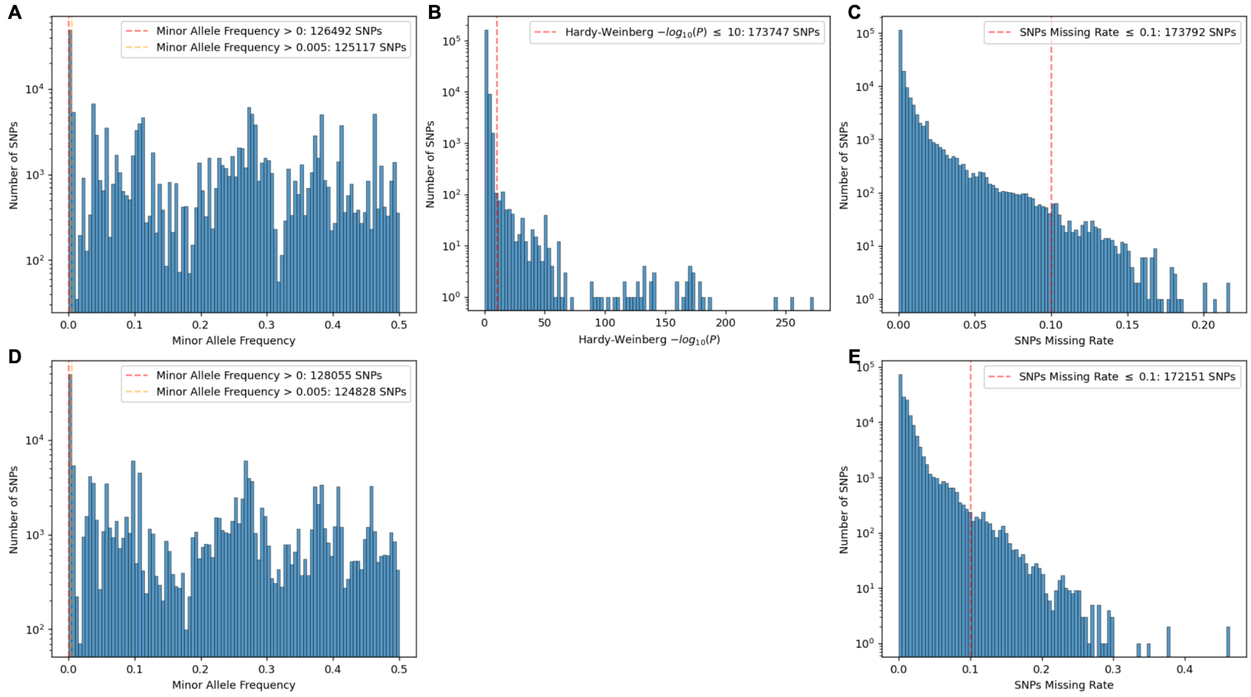

**Supplementary Figure S7.** Mitochondria SNPs statistics. **A.** Mitochondria SNPs minor allele frequency histogram. **B.** Mitochondria SNPs missing rate histogram.

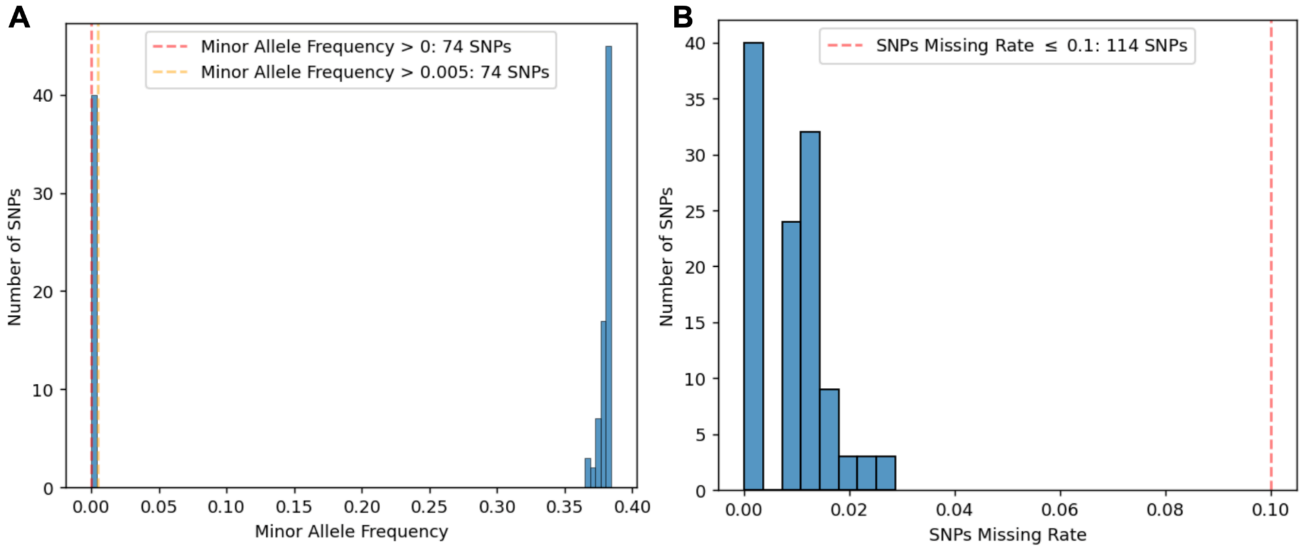

**Supplementary Figure S8.** Principal component analysis on genotypes produced using different sequencing methods. PCs that explain more than 10% variance are shown.

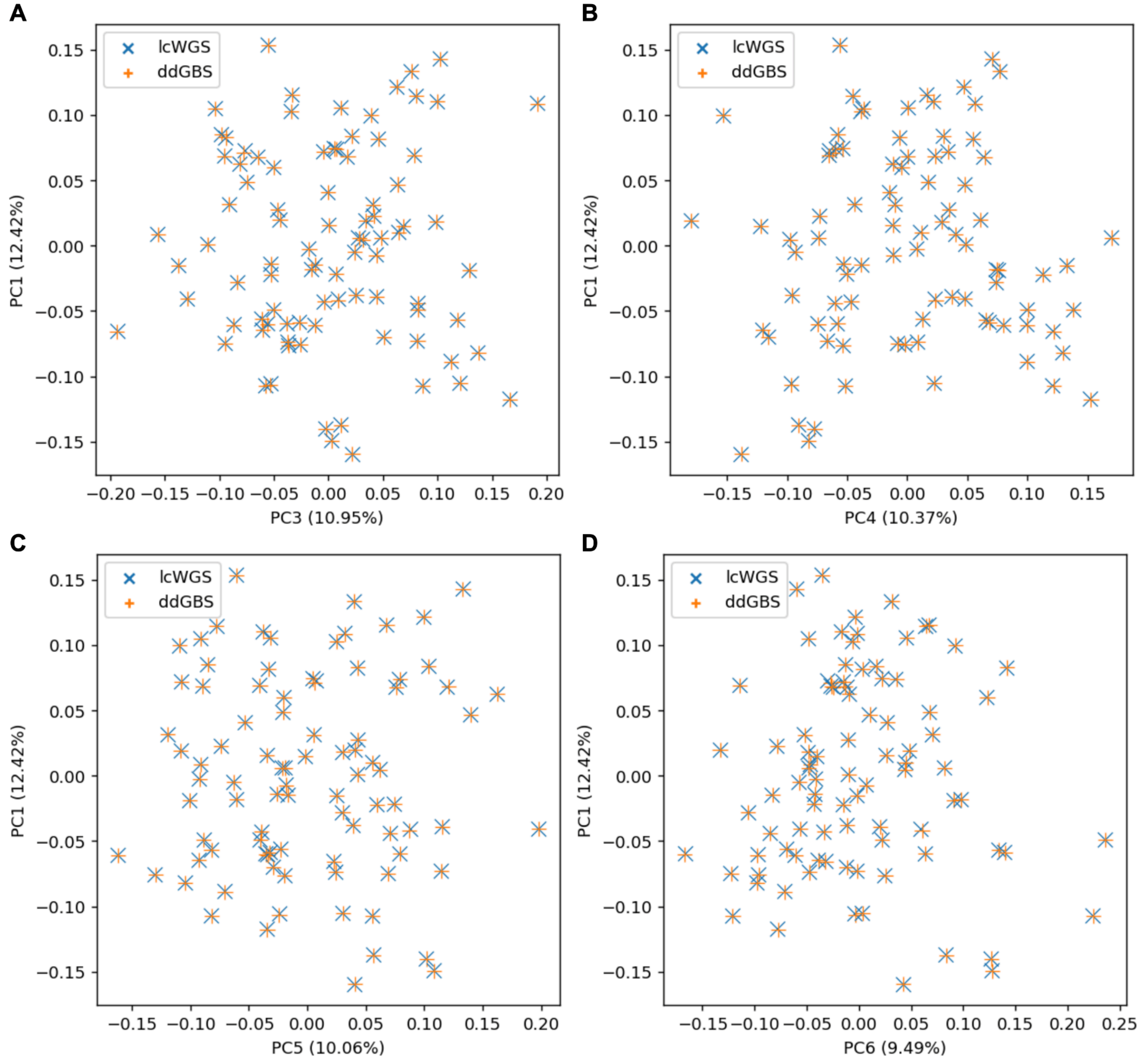

### References

Bushnell B. BBTools. http://sourceforge.net/projects/bbmap/.

Danecek P, Bonfield JK, Liddle J, Marshall J, Ohan V, Pollard MO, Whitwham A, Keane T, McCarthy SA, Davies RM, et al. 2021. Twelve years of SAMtools and BCFtools. GigaScience. 10(2):giab008. doi:10.1093/gigascience/giab008.

Li H. 2013. Aligning sequence reads, clone sequences and assembly contigs with BWA-MEM. [accessed 2023 Apr 10]. http://arxiv.org/abs/1303.3997.

Poplin R, Chang P-C, Alexander D, Schwartz S, Colthurst T, Ku A, Newburger D, Dijamco J, Nguyen N, Afshar PT, et al. 2018. A universal SNP and small-indel variant caller using deep neural networks. Nat Biotechnol. 36(10):983–987. doi:10.1038/nbt.4235.

Van der Auwera GA, O'Connor BD. 2020. Genomics in the Cloud: Using Docker, GATK, and WDL in Terra. 1st ed. O'Reilly Media.

Yun T, Li H, Chang P-C, Lin MF, Carroll A, McLean CY. 2021. Accurate, scalable cohort variant calls using DeepVariant and GLnexus. Robinson P, editor. Bioinformatics. 36(24):5582–5589. doi:10.1093/bioinformatics/btaa1081.
